# *Escherichia coli* chemotaxis to competing stimuli in a microfluidic device with a constant gradient

**DOI:** 10.1101/2021.12.30.474376

**Authors:** Xueying Zhao, Roseanne M. Ford

**Affiliations:** Department of Chemical Engineering, University of Virginia, Charlottesville, Virginia

**Author notes:** **Correspondence**, Roseanne M. Ford, Department of Chemical Engineering, University of Virginia, 102 Engineers’ Way, Charlottesville, VA 22904.

**Keywords:** bacterial chemotactic response, two-component signaling system, multiscale modeling, multiple stimuli, microfluidic device, constant gradient chamber

## Abstract

In natural systems bacteria are exposed to many chemical stimulants; some attract chemotactic bacteria as they promote survival, while others repel bacteria because they inhibit survival. When faced with a mixture of chemoeffectors, it is not obvious which direction the population will migrate. Predicting this direction requires an understanding of how bacteria process information about their surroundings. We used a multiscale mathematical model to relate molecular level details of their two-component signaling system to the probability that an individual cell changes its swimming direction to the chemotactic velocity of a bacterial population. We used a microfluidic device designed to maintain a constant chemical gradient to compare model predictions to experimental observations. We obtained parameter values for the multiscale model of *Escherichia coli* chemotaxis to individual stimuli, α-methylaspartate and nickel ion, separately. Then without any additional fitting parameters, we predicted the response to chemoeffector mixtures. Migration of *E. coli* toward α-methylaspartate was modulated by adding increasing concentrations of nickel ion. Thus, the migration direction was controlled by the relative concentrations of competing chemoeffectors in a predictable way. This study demonstrated the utility of a multiscale model to predict the migration direction of bacteria in the presence of competing chemoeffectors.

## 1. Introduction

Bacteria are exposed to a milieu of chemical stimulants in their natural surroundings. In agricultural soils microbes fix nitrogen in the presence of root exudates (Bergman, 1990; Anderson et al., 1993; Babalola, 2010), in polluted groundwater aquifers microorganisms degrade hydrocarbon mixtures (Pandey and Jain, 2002; Aburto and Peimbert, 2011; Meckenstock et al., 2015), and in the gut microbiome bacteria are exposed to a wide array of nutrients and toxins (Flint et al., 2012; Keilberg and Ottemann, 2016; Feng et al., 2019; Tsiaoussis et al., 2019). Facing a specific chemical gradient, a population of bacteria can direct their movement to swim toward or away from it, exhibiting a behavior termed chemotaxis. In some chemotactic responses, bacteria are attracted to a higher chemical concentration because they can use these chemicals as food and energy sources. Such compounds are called chemoattractants, while compounds bacteria swim away from are called chemorepellents. For example, *Escherichia coli* show positive chemotaxis to amino acids and aromatic compounds, while sulfides and inorganic ions cause bacteria to exhibit negative chemotaxis (Terracciano et al., 1984, Pandey and Jain, 2002).

The physical swimming mechanism for chemotaxis of *E. coli* is well known (*e.g*. Sourjik, 2004); when the flagella rotate clockwise (CW), bacteria will tumble in place, while the counterclockwise (CCW) rotation of the flagella will lead to a “run” along a straight pathway. In the absence of chemoeffectors, bacteria perform a random walk similar to Brownian motion (Rivero et al., 1989) alternating between runs of ~1s and tumbles of ~0.1 s (Berg and Brown, 1972). In the presence of a chemoattractant gradient, bacteria extend their run times toward higher chemoattractant concentrations by decreasing the tumble probability. This results in the bacterial population showing a net drift velocity toward the chemoattractant. In contrast, in the presence of a chemorepellent gradient, cells increase their tumbling frequency when moving in an increasing chemorepellent concentration (Tso and Adler, 1974). Thereby, the bacterial population exhibit a net drift velocity down the chemorepellent gradient.

Most previous studies measured bacteria chemotaxis in the presence of only one chemoeffector. However, bacteria encounter multiple chemicals in their native environments. Thus, it is important to investigate how bacteria respond to conflicting information from multiple chemical signals. Several researchers have studied bacterial chemotactic responses in the presence of multiple stimuli. Mowbray and Koshland (1987) studied the interaction of aspartate and maltose stimuli with the Tar receptor. They reported that the response was additive and independent because aspartate and maltose bind to separate sites on the Tar receptor. However, Strauss et al. (1995) argued that the response of fucose and α-methylaspartate were not addictive because the simple model predictions did not match the experimental results they performed with multiple stimuli and proposed that a more complex relationship which incorporated signal processing steps was required. Zhang et al. (2019) observed that a traveling escape band formed in opposing gradients of aspartate and tryptone broth. This phenomenon was explained by bacteria first responding to a strong attractant, that was also a poor nutrient (aspartate) at one end of the channel. Then subsequently, the consumption of nutrient resulted in bacteria traveling toward a rich nutrient, that was also a weak attractant (tryptone broth) at the other end of the channel. Kalinin et al. (2010) found that the ratio of two receptors (Tar and Tsr) determined bacteria chemotactic response to α-methylaspartate (MeAsp) and serine. Their study indicated when the ratio of receptors Tar/Tsr is more than a threshold of 2, bacteria prefer to swim toward serine instead of MeAsp. Park and Aminzare (2022) used mathematical modeling and found the agreement with the experimental results in Kalinin et al. (2010). However, most of these studies measured the behavior of a population of bacteria, without emphasizing how the signal transduction mechanism functions inside an individual bacterium. In order to predict the response to multiple stimuli, it is important to relate the cellular-level signal transduction mechanism to population-scale chemotaxis behavior. By using multi-scale modeling, we can link molecular reaction to individual cell level swimming behavior, and then relate that to population-scale dynamics. In this paper, we quantified chemotaxis using a mathematical model (Middlebrooks et al., 2021) that integrates signal transduction kinetics from multiple inputs, starting from chemoeffector binding to chemotactic receptor, to the run and tumble behavior of individual cells, and then to the population-scale chemotactic velocity.

At the molecular scale, the signaling response in *E. coli* chemotaxis depends on the phosphotransfer between a histidine kinase and a response regulator (Sourjik et al., 2010). The binding of chemoattractant inactivates the kinase and decreases the tumble probability, while the binding of chemorepellent increases the activity of the kinase and increases tumble probability (Grebe and Stock, 1998; Jasuja et al., 1999). This change of tumble probability can further relate to bacteria response at the individual cell level since the mean run time is the reciprocal of the tumbling probability. The individual scale response is developed from a theoretical model describing chemotactic behavior by Rivero et al. (1989). They used the empirical correlation developed by Berg and Brown (1972), in which the logarithm of mean run time increases with respect to the rate of change in the number of bound receptors, which is related to the change in concentration of signaling complexes inside the cell.

To facilitate a quantitative measure of the chemotactic response at the population scale, it is beneficial to create a constant chemical gradient, which will serve as the driving force for chemotaxis. Middlebrooks et al. (2021) used a stopped flow diffusion chamber to analyze bacterial chemotactic response to the combination of aspartate and nickel ion. The device they used was able to create a sharp step change in the chemical concentration of chemoeffector that relaxed over time in a predictable way due to diffusion. However, the design did not maintain a constant chemical gradient during bacterial chemotactic response. In comparison, the microfluidic device used by Wang et al. (2015) provided more easily quantifiable results since it was able to reach a steady state condition with a constant chemical concentration gradient. We adapted this unique microfluidic device (Wang et al, 2015) to produce a constant concentration gradient in the channel that is also free of fluid convection.

We exposed bacteria to different chemicals (including attractants, repellents and their combination) with constant gradients using the microfluidic design of Wang et al. (2015). We used *E. coli* to study bacterial chemotaxis with the presence of conflicting stimuli α-methylaspartate and nickel ion. Experimental data were used to obtain parameters in a mathematical model capturing bacterial motility and chemotaxis. We incorporated parameters from single-stimulus responses into our multiple stimuli model to gain a better picture of bacterial chemotaxis in a heterogeneous chemical environment.

## 2. Mathematical Model

### 2.1 Bacteria transport processes at the population level

Bacterial transport in the cross channels on the bottom layer of the microfluidic device (Figure 1) was modeled using species mass conservation equations. A one-dimensional equation was deemed suitable following the work of Wang et al. (2015). They observed no dispersion in the cross channel for a tracer molecule. Thus, mass transfer between the vias through the cross channel was controlled by diffusion only; the convection term was eliminated from the equation as the device design prevents bulk fluid flow in the cross channel.

**Figure 1.**
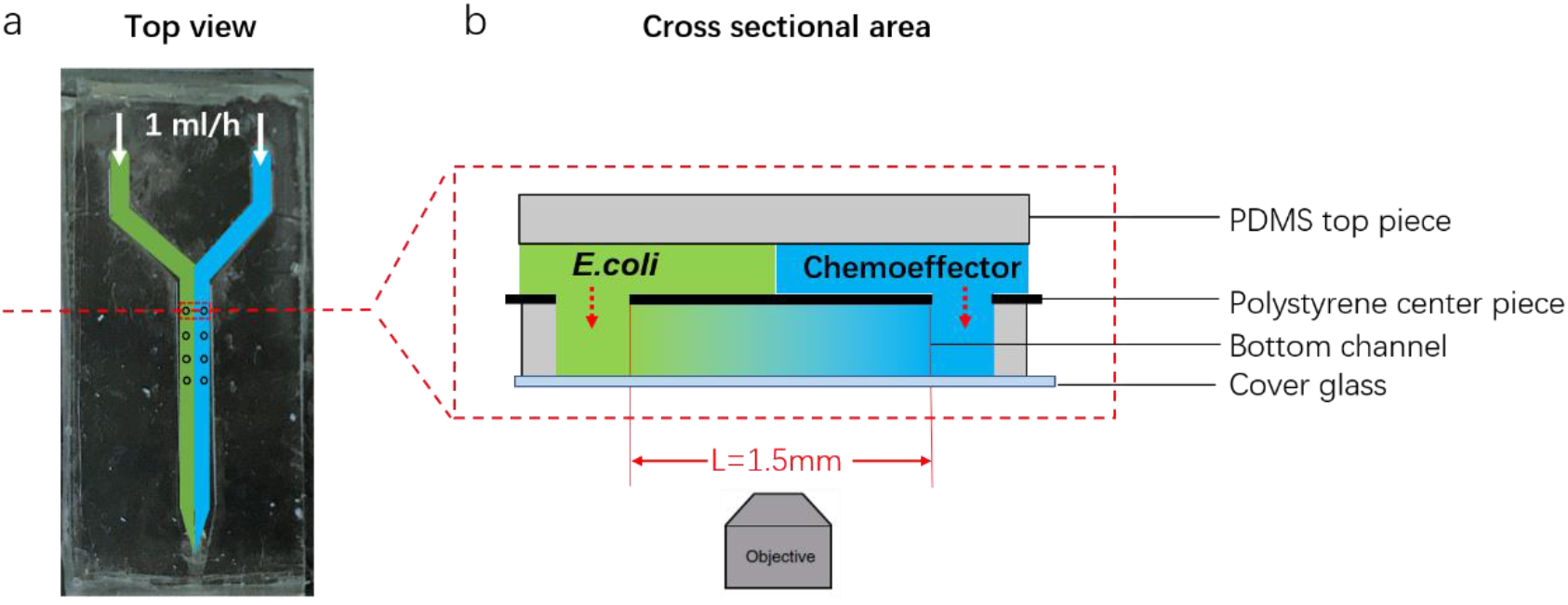
(a) Photo of microfluidic device (top-down view) with *E. coli* flow path colored green and chemoeffector colored blue and (b) an enlarged cross-sectional view showing connections between the top and bottom channels (not drawn to scale in order to emphasize gradient formed by diffusion in the cross channel).

In the absence of chemoeffectors, the governing equation for bacteria concentration *b* under unsteady state conditions is

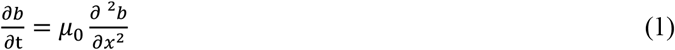

where *μ*_0_ is the bacterial random motility coefficient. Note that *μ*_0_ can only be evaluated under unsteady state conditions as it will cancel out at steady state and the bacterial distribution becomes independent of *μ*_0_. The boundary conditions are *b* (0, *t*) = *b*_0_, *b* (*L, t*) = 0 and the initial condition is *b* (*x*, 0) = 0. The solution to Equation (1) is

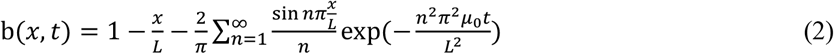

In the presence of a chemoeffector, the governing equation for bacteria concentration *b* at steady state conditions is

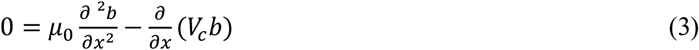

where *V_c_* is the chemotactic velocity as derived in Middlebrooks *et al*. (2021) for multiple chemoeffectors. The equations for *V_c_* under different conditions (attractant, repellent and their combination) are shown in the Supplementary Information. The bacteria concentration *b* was solved using finite difference method in MATLAB R2018b (MathWorks).

Chemotactic velocity in the presence of attractant is

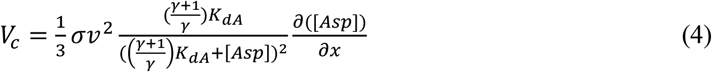

where *σ* is the stimuli sensitivity coefficient, *v* is bacteria swimming speed, *γ* is the signaling efficiency, [*Asp*] is attractant concentration, *K_dA_* is the dissociation constant of attractant binding. The equations for chemotactic velocity *V_c_* under other cases and the derivation are provided in the Supplementary Information.

The governing equation for chemoeffectors at steady state is

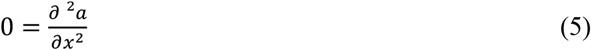

where *a* is the chemoeffector concentration, and *x* is the position in the cross channel. *E. coli* do not metabolize α-methylaspartate or nickel, so a consumption term was not needed in the governing equation. The boundary conditions are *a* (x=0) =0 and *a* (x=*L*) = *a*_0_, where *L* is the length of the channel. The solution of the equation is a linear concentration profile with a constant gradient (d*a*/dx = *a*_0_/L). The chemoeffector concentration and gradient are needed to evaluate the chemotactic velocity *V_c_* term in Equation 3.

MATLAB R2018b was used to solve the equations, plot the bacterial concentration distribution in the cross channel, and fit the parameters. The *μ*_0_ value was obtained by using least square regression analysis to fit Equation 2 to experimental data from unsteady state conditions without addition of chemoeffector. Equation 3 was solved using the chemoeffector concentration and gradient (from Equation 5) to get bacterial distributions given set of chemotaxis parameter values and then plot those distributions. Chemotactic parameters values were obtained by using least square regression analysis to fit Equation 3 to experimental data from steady state conditions in the presence of chemoeffectors.

### 2.2 Two-component signaling kinetics for chemotaxis

In the absence of chemoattractant, the kinase CheA has autophosphorylation activity. The CheA phosphoryl group is further transferred to response regulator CheY as depicted in Figure 2a. The response regulator diffuses through the cytoplasm and transmits the signal to a flagellar motor, enhancing the probability of CW rotation and causing bacteria to tumble (Sourjik et al., 2010).

**Figure 2.**
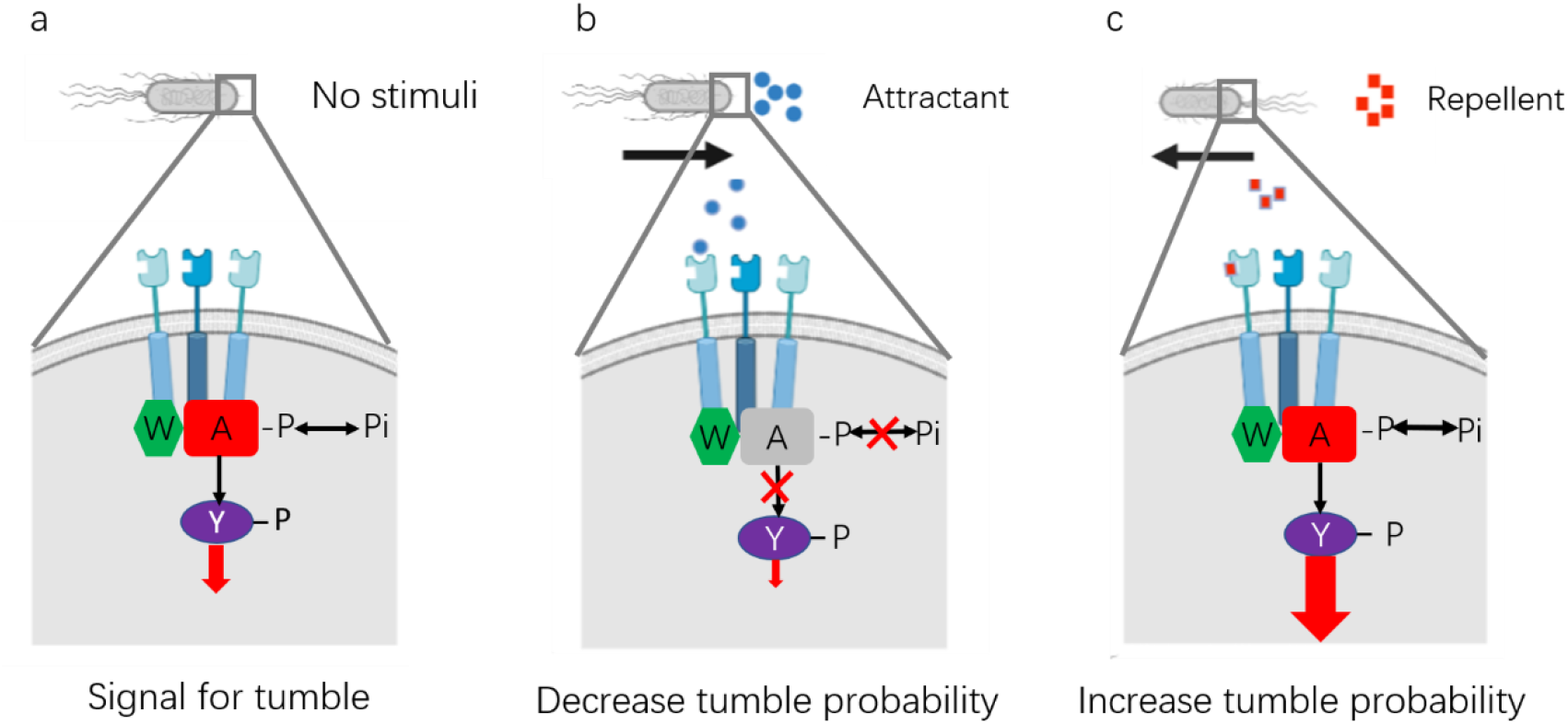
Schematic of the two-component signaling system in *E. coli* (a) in the absence of chemical stimuli, (b) when the Tar receptor complex is bound by attractant, and (c) when the Tar receptor complex is bound by repellent. A, W, and Y indicate chemotaxis proteins CheA, CheW and CheY respectively; P is phosphate. Adapted from Sourjik and Armitage (2010).

In the presence of a chemoattractant, binding of the attractant causes CheA inactivation, decreases the probability of clockwise (CW) rotation and causes the bacterium to tumble less frequently, which leads to the bacterium continuing to move in the same direction. On the other hand, the binding of repellent does not inactivate CheA, and instead appears to increase the CheA activity, which increases the probability of clockwise (CW) rotation leading to more frequent tumbles and bacteria tending to swim away from the repellent source.

Detailed equations for the multiscale model are shown in the Supplementary Information. Three parameters were used in the model. The stimuli sensitivity coefficient *σ* captures the ratio of change in tumbling frequency to the change in the concentration of signaling complex [*CheA* − *P*], and is only specific to the bacterium. The signaling efficiency *γ* represents the ratio of dissociation constant of [*CheA*] phosphorylation to the concentration of phosphate, which is essentially the ratio of [*CheA*] to [*CheA* − *P*]. The parameter *γ* can be seen as the inverse of the gain in the signaling process. The repellent sensitivity coefficient *κ* represents the ratio of dissociation constant of phosphorylation reaction of [*CheA*] to the dissociation constant of phosphorylation reaction of repellent bound [*CheA*], and is specific to the repellent. *κ* represents the enhancement of repellent bound receptor in the phosphorylation signaling process compared to that in the absence of chemoeffector.

## 3. Experimental Methods

### 3.1 Preparation of bacteria cultures

*E. coli* HCB1 (Wolfe et al., 1987) were previously transformed to express green fluorescent protein (GFP). The transformation was performed using the protocols from the pGLO Bacterial Transformation Kit (Bio-Rad Laboratories, Hercules, CA). In each experiment, 100 μL of GFP-labeled *E. coli* HCB1 frozen stock was cultured in 50 mL Luria broth (Fisher Scientific, NY) in a 250 mL Erlenmeyer flask on a Thermo Scientific Incubated Shaker (MaxQ4000) with a rotation rate of 150 rpm at 30 °C. 10 μL ampicillin (100mg/mL, Sigma, MO) and 1 mL 100mg/mL D(-)-arabinose (Fisher Scientific, NY) were added into growth media to provide selective pressure for maintaining the plasmid and expressing GFP.

Bacteria were harvested at the mid-exponential phase of population growth and resuspended in the random motility buffer (RMB) to reach an optical density of 1.20 at 590 nm (measured in spectrophotometer, Molecular Devices, Spectramax 384 plus). Mid-exponential phase corresponds to the conditions when bacteria exhibit the greatest motility (Worku et al., 1999). Then bacteria were filtered from the culture media with 0.22 μm membrane filter (Millipore, MA) using vacuum filtration (Berg and Turner, 1990) and resuspended into 5% RMB (RMB, including 9.13 g/L Na_2_HPO_4_ (Fisher Scientific), 4.87 g/L NaH_2_PO_4_·H_2_O (Amresco), and 0.029 g/L EDTA (Sigma, MO)) to an optical density around 1.0 (8 × 10^8^ cells/mL). Before performing motility and chemotaxis experiments, the motility of the bacteria was examined under a Zeiss 100/1.25 oil lens with a Nikon microscope (Digital Sight DS-5Mc).

### 3.2 Microfluidic design, fabrication and operation

The uniquely designed microfluidic device shown in Figure 2 is made up of three layers (Wang et al., 2015). Both the top and bottom channels are made of polydimethylsiloxane (PDMS). PDMS was chosen because of its oxygen permeability, allowing the bacteria to maintain motility inside the channel. The detailed steps of making PDMS pieces are described in the Supplementary Information. The top layer has two inlets connected to the main channel and the dimensions of the main channel are 95 μm high, 3.3 mm wide, and 2 cm long. The centerpiece that fits between the two PDMS layers was made from black polystyrene material (Staples Inc.). The purpose of the black color is to block background fluorescence in the main channel from the signal in the bottom cross-channels. Double-sided tape (3M, MN) was used to adhere the centerpiece to the top and bottom layers. As depicted in Figure 2 four pairs of vias were cut into the centerpiece using a laser cutter (VersaLaser, AZ). The vias connect the main channel in the top layer with the cross channels buried in the bottom layer. Because the pressures are equal in the vias at the cross positions perpendicular to the flow direction, no convection flow will occur in the bottom channels; there is only diffusion in the bottom channel. The dimensions for the cross channels were 100 μm high, 600 μm wide, and 1.5 mm long.

The bacteria suspension and chemoattractant (or 5% RMB for control experiments) were introduced into opposite arms leading to the main channel. A syringe pump SP 220i (World Precision Instruments, LLC, FL) was used to control flow at a constant volumetric flow rate of 1.0 mL/h, which corresponds to a linear speed of 2.42 mm/s. The Reynolds number in the main channel with the connecting vias was 0.17, which means the two streams were under laminar flow.

### 3.3 Microscopy and image analysis

The fluorescence images were recorded using a Zeiss 780 confocal microscope with 10x objective lens in the Keck Center for Cellular Imaging at the University of Virginia. To observe the signal from bacteria, a MBS 488/561 filter was used and the detector was in the range of 502-607 nm. Images were collected at the speed of 1.94 seconds per frame, which gave us a 5.25 MB image. The 1.5 mm-long cross channel was taken in two images, and then stitched together with overlap percentage 0.01%. Bacterial intensity was recalibrated by automatic adjustment of the lighting as the microscope camera takes image tiles along the channel; this recalibration is programmed into the imaging software. Five sequential images were collected for each region and superimposed together. For each image, the region was selected starting from the edge of the bacteria source via. The pixel between scale markings was measured from the image and then scaled to the known distance of 0.4 mm from the design. Then the red rectangular box was drawn to select the analysis region. The gray level value in each region was collected by using the plot profile feature.

The following equation was used to normalize the fluorescence intensity for each image:

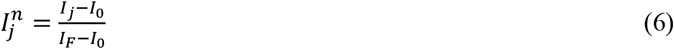

where *I* represents the gray level value, the superscript n represents the normalized value, subscript j corresponds to certain location along the channel, *I*_0_ is the gray level value at the bacteria sink, *I_F_* is the gray level value at the bacteria source.

## 4. Results and Discussion

The random motility coefficient of the bacterial population was evaluated in the absence of chemoeffectors under unsteady state conditions. Then, attractant only and repellent only experiments were performed under steady state with constant chemoeffector gradients to obtain chemotactic parameters for each individual stimulus. These parameters were incorporated into the multiscale model to predict bacterial distributions when exposed to a combination of chemoeffectors and compared to experimental observations.

### 4.1 *E. coli* HCB1 random motility coefficient

To evaluate the random motility coefficient (*i.e*. bacterial diffusivity), a constant source of GFP-labeled chemotactic bacteria *E. coli* HCB1 was maintained at one end of the cross channel. The distribution of bacteria as they migrated through the cross channel is shown in Figure 3a at several times; the corresponding normalized bacteria fluorescence intensity is plotted in Figure 3b. The qualitative shape of the bacterial profiles followed the expected exponential decay in concentration from the constant source at x=0 as well as the increasing degree of penetration into the cross channel over time. At long times the normalized bacterial distribution approached a decreasing linear trend as expected once steady state was achieved (~ 60 min). A best-fit of the mathematical model (Equation 2, n=10) to experimental data using nonlinear regression yielded a random motility coefficient of *μ*_0_ = 1.3 ± 0.21 × 10^−10^ *m*^2^/*s* (averaged over four replicates). This motility coefficient value was similar in magnitude to others that were previously reported (*e.g* 2.4 × 10^−10^ *m*^2^/*s* for *E. coli* in Wang *et al*., 2015). Theoretical predictions using the experimentally-derived value for *μ*_0_ in Equation 2 are plotted in Figure 3b for comparison to experimental observations. We also note that this population scale parameter, the random motility coefficient, can be calculated from individual cell swimming properties such as the tumbling probability, which depends on molecular levels of phosphorylated CheY and the signaling efficiency parameter *γ*. Thus, the measurement of the random motility coefficient provides an independent means for estimating the value of the signaling efficiency parameter. Details of this analysis are included in the Supplementary Information.

**Figure 3.**
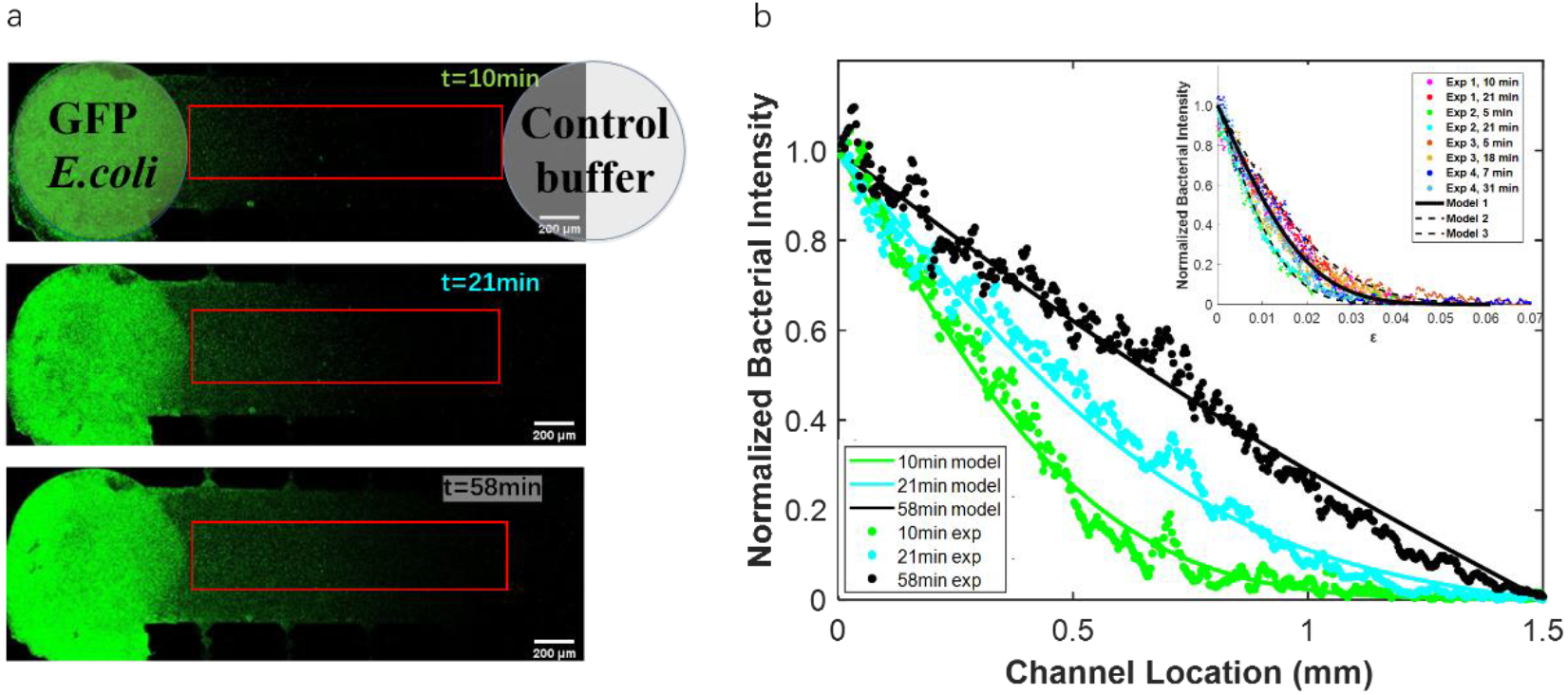
(a) GFP-labeled *E. coli* distribution in a cross channel at several times. There is a constant source of bacteria on the left-hand side of the channel and a constant sink on the right-hand side; the fluorescence intensity is proportional to bacterial concentration. Red rectangular boxes indicate the region over which data was analyzed. (b) Normalized bacterial intensity along the channel (scattered data) compared with model outcomes (solid lines) at several times. The random motility coefficient obtained from this set of data was *μ*_0_ = 1.6 × 10^−10^ *m*^2^/*s*. (*Inset*) Normalized bacteria intensity with respect to the Boltzmann transformation parameter for four sets of experimental data collected at unsteady state, where 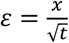. Model 1 corresponds to the solid curve with *μ*_0_ = 1.3 × 10^−10^ *m*^2^/*s*, Model 2 corresponds to the upper dashed curve with *μ*_0_ = 2.1 × 10^−10^ *m*^2^/*s*, and Model 3 corresponds to the lower dashed curve with *μ*_0_ = 0.70 × 10^−10^ *m*^2^/*s*.

For unsteady diffusive processes the Boltzmann transformation combines the spatial and temporal dependence into a single parameter according to the equation

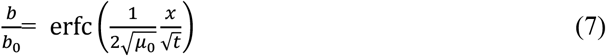

where 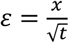 with units of 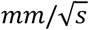 This transformation faciliates comparison of experimental results and model predictions at multiple time points. The normalized bacteria fluorescence intensity when plotted with respect to the transformation paramete *ε* yielded a single curve as shown in Figure 3b Inset, confirming that bacterial migration followed a diffusive process with *μ*_0_ = 1.3 × 10^−10^ *m*^2^/*s*.

### 4.2 *E. coli* HCB1 chemotactic response to an attractant *α*-methylaspartate

A constant source of GFP-labeled chemotactic bacteria *E. coli* HCB1 was provided at one end of the cross channel, while a constant source of the attractant 0.2 mM *α*-methylaspartate was provided at the opposite end of the channel. The distribution of bacteria in the cross channel after reaching steady state (~ two hours) or longer is shown in Figure 4a; the normalized bacteria intensity is shown in Figure 4b. The distribution of chemotactic bacteria followed a curved parabolic shape with positive deviation from the control case (without chemoeffector) because of increased migration due to chemotaxis. A multiscale mathematical model for bacterial transport with chemotaxis (solved numerically with MATLAB software, The MathWorks, Inc.) was used to generate the bacterial distribution profile by solving Equation 3 with the chemotactic velocity as specified for *α*-methylaspartate in Equation 4. Experimental data collected from bacteria responses to two different concentrations of attractant (0.2 mM and 0.4 mM) were used to obtain the values of *σ* and *γ*. A non-linear least squares algorithm was used to regress the parameters in the theoretical model to align with experimental data. The set of parameter values used in the model are shown in Table 1. The parameter values are consistent with others that have been previously reported (Middlebrooks et al., 2021; Middlebrooks, 1993; Clarke and Koshland, 1979). From Equation 4 for the chemotactic velocity, the maximum chemotactic velocity occurs when the concentration of chemoattractant is closest to the *K_dA_* value. This explains the greater overall response of bacteria to 0.4 mM *α*-methylaspartate (Figure 5c) compared to 0.2 mM *α*-methylaspartate because *K_dA_*=0.64 mM.

**Figure 4.**
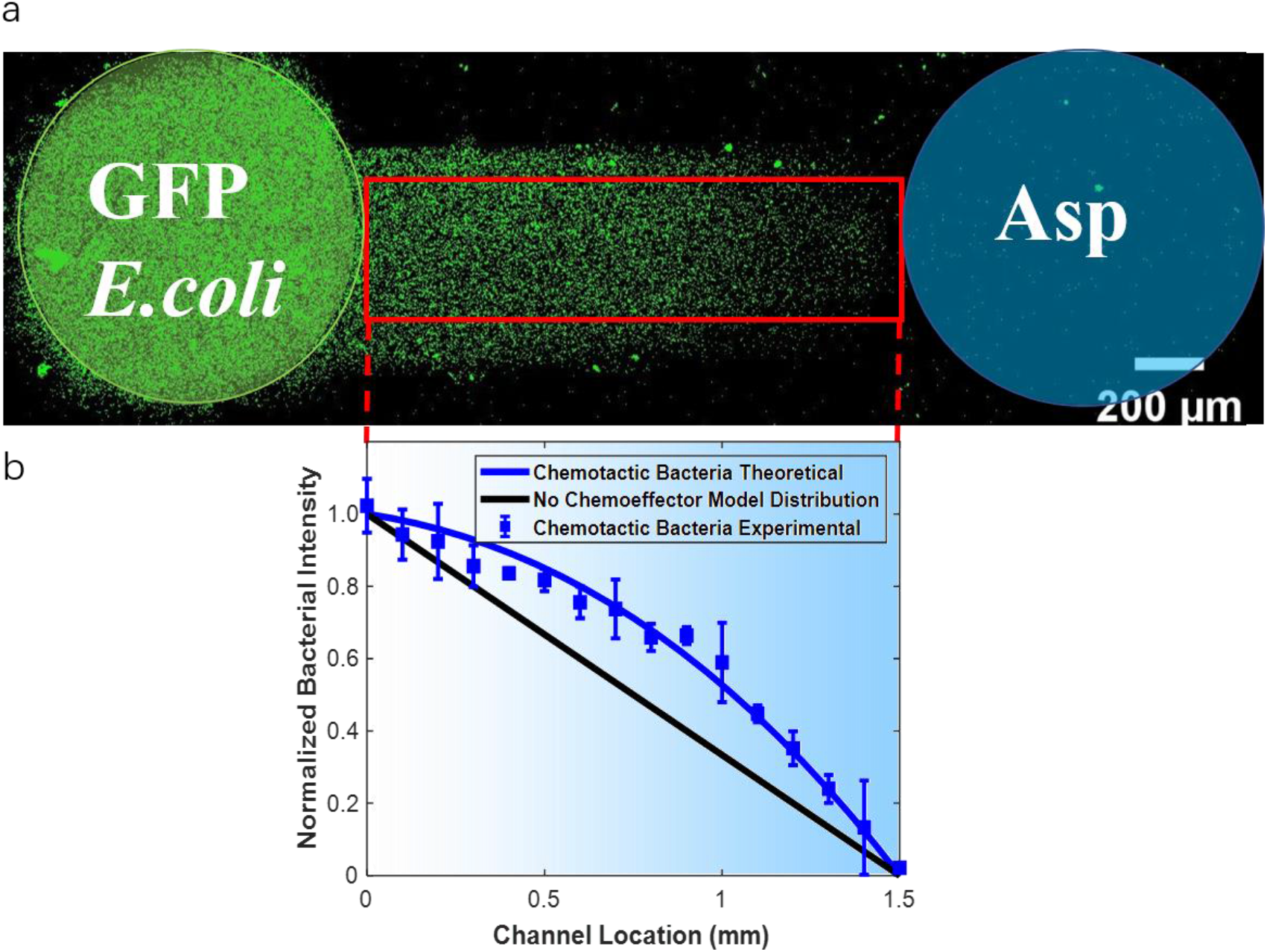
(a) GFP-labeled *E. coli* distribution in a cross channel at steady state. There is a constant source of bacteria on the left-hand side of the channel, and a constant attractant source of 0.2 mM *α*-methylaspartate on the right-hand side; the fluorescence intensity is proportional to bacterial concentration. The red rectangular box indicates the region over which data was analyzed. (b) Normalized bacteria intensity along the channel (scattered data) and the model results (solid lines). The black line indicates the expected bacterial distribution at steady state for the control case without chemoattractant. The blue shading from right to left represents the decreasing attractant concentration.

**Figure 5.**
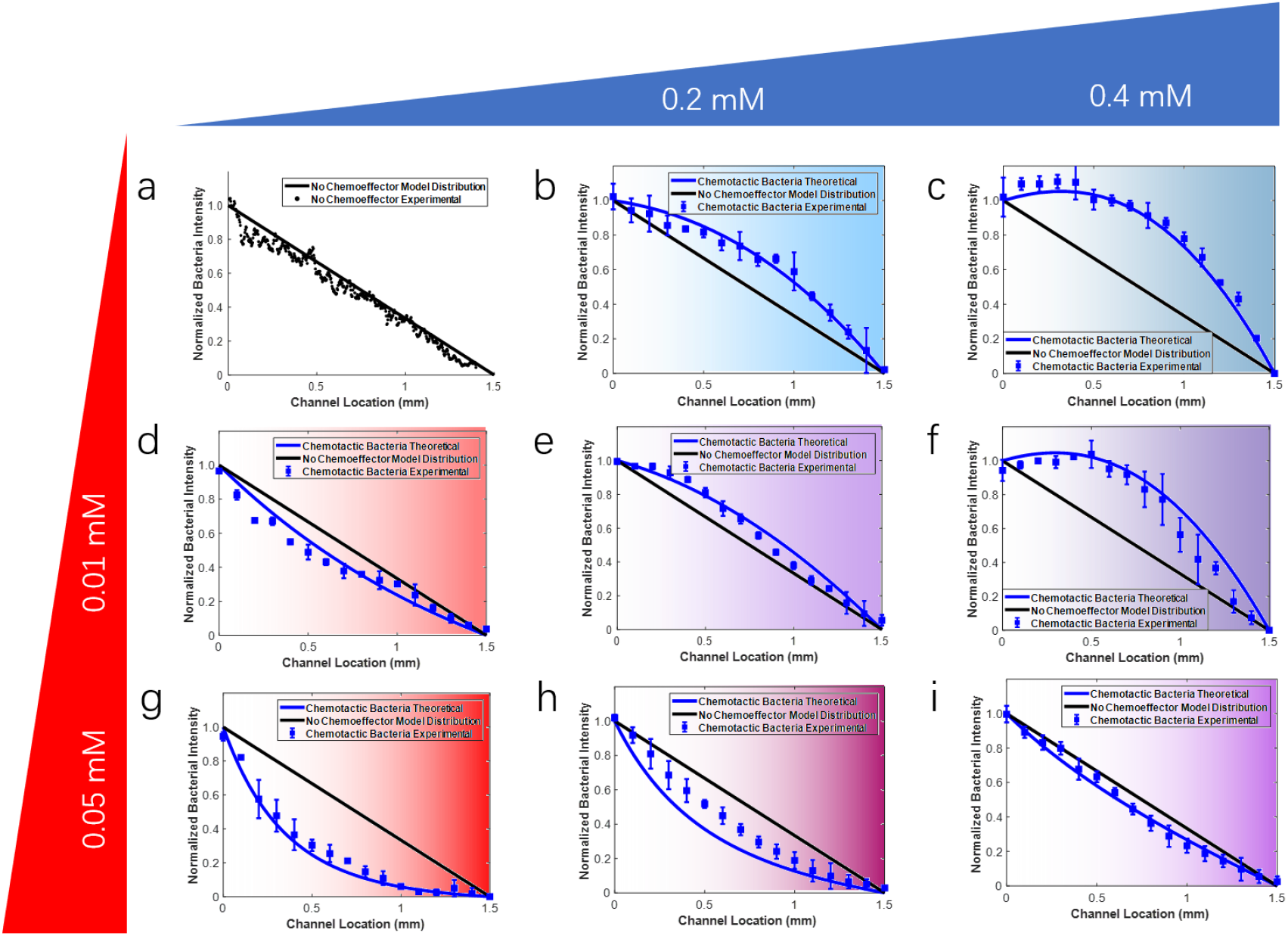
GFP-labeled *E. coli* distributions in a cross channel at steady state. There is a constant source of bacteria on the left-hand side of the channel and a constant source of buffer, attractant and/or repellent on the right-hand side: (a) buffer only as a control, (b) 0.2 mM attractant *α*-methyl aspartate, (c) 0.4 mM attractant, (d-f) a combination of 0.01 mM repellent nickel ions and 0, 0.2 or 0.4 mM attractant, respectively, (g-i) a combination of 0.05 mM repellent and 0, 0.2 or 0.4 mM attractant, respectively. The fluorescence intensity is proportional to bacterial concentration.

**Table 1.** Parameters used in the model and comparison with reference values

| Symbol | Parameter | Model Value | Ref Value |
| --- | --- | --- | --- |
| $\gamma$ | Signaling efficiency | 4 | 20 (Middlebrooks et al., 2021) |
| $\sigma$ | Stimuli sensitivity coefficient (s) | 9 | 11 (Middlebrooks et al., 2021) |
| $\kappa$ | Repellent sensitivity coefficient | 10 | 14 (Middlebrooks et al., 2021) |
| $v$ | Swimming speed ($\mu\text{m/s}$) | 22 | 22 (Wang, 2013) |
| $K_{dA}$ | Dissociation constant for α-methyl aspartate binding (mM) | 0.64 | 0.64 (Clarke et al, 1979) |
| $K_{dN}$ | Dissociation constant for nickel ion binding (mM) | 0.54 | 0.54 (Middlebrooks et al., 2021) |
| $\mu_0$ | Random Motility Coefficient ($\text{m}^2/\text{s}$) | $1.3 \times 10^{-10}$ | $2.4 \times 10^{-10}$ (Wang, 2013) |

### 4.3 *E. coli* HCB1 chemotactic response to a repellent nickel ion

We measured the response of GFP-labeled *E. coli* HCB1 to repellent nickel ions in the cross channel. Bacteria were provided as a constant source at one end of the channel and the repellent, either 0.01mM or 0.05 mM nickel sulfate, was provided as a constant source at the opposite end of the channel. Distributions of bacteria after reaching steady state (~2 hrs) are shown in Figure 5d and Figure 5g. They show a curved parabolic shape with a negative deviation from the control case because of chemotactic migration away from the nickel source. A greater negative response of bacteria to 0.05 mM nickel (Figure 5g) over 0.01 mM nickel (Figure 5d) was observed because the higher concentration is closer to *K_dN_*=0.54 mM, which represents the nickel ion concentration to which cells are most responsive. A mathematical model for bacterial transport with chemotaxis (solved numerically with MATLAB R2018b, The MathWorks, Inc.) was used to predict the bacterial distribution profile by solving Equation 3 and using the chemotactic velocity given by Equation S24 for nickel ion. The model was solved with values of *σ* and *γ* obtained from attractant case and then fit to data from the repellent experiments to obtain a value for *κ*.

### 4.4 *E. coli* response to chemoeffector mixtures

We measured the response of GFP-labeled *E. coli* HCB1 to different combinations of attractant *α*-methylaspartate and repellent nickel ion in the cross channel. There was a constant source of bacteria on the left-hand side of the channel. Different combinations of attractant *α*-methylaspartate (0.2 and 0.4 mM) and repellent nickel sulfate (0.01 and 0.05 mM) were provided as a constant source of chemoeffectors on the right-hand side. In the presence of both attractant and repellent, the normalized population density of bacteria in the channel fell somewhere between the distributions for the attractant only and repellent only cases, which indicates that bacteria integrate chemical information taking into account both chemoeffectors. The bacterial profile in Figure 5e shows a curved parabolic shape with positive deviation from the control case when bacteria responded to the combination of 0.2 mM *α*-methylaspartate and 0.01 mM nickel sulfate, On the contrary, as the nickel sulfate concentration increased to 0.05 mM in Figure 5h we observed a negative deviation from the control case. The negative deviation was due to the repellent response overwhelming the attractant response for that particular combination. As the *α*-methylaspartate concentration was increased to 0.4 mM in Figure 5i, we still observed a net negative deviation from the control case. However, the increased attractant concentration countered the repellent effect and moved the bacterial distribution closer to the control case, as the effect of attractant and repellent almost canceled out each other for this concentration pairing.

MATLAB software (The MathWorks, Inc.) was used to predict the bacterial distribution profile by solving the governing Equation 3 for bacteria and using the chemotactic velocity of Equation S26 for the combined cases. We applied the parameters in Table 1 without further fitting to obtain the theoretical predictions in the combined cases. The model predicted the overall direction of bacterial migration correctly and did a good job capturing the experimental bacterial distributions for the mixtures. In one case (Figure 5e) the model predicted a greater repellent effect than was observed in the experimental data. Because the parameters were obtained by fitting the experimental results for the attractant and repellent only cases, it is not surprising that the model is more likely to match responses where one chemical dominates (Figure 5f and Figure 5h), while less predictive when the responses almost cancel out (Figure 5e and Figure 5i).

The results summarized in Figure 5 show that the multiscale model and its parameters, capturing bacteria chemotaxis signal transduction steps inside the cell, provided a good description of bacterial population behavior. Thus, the multiscale model can be used to predict bacterial chemotaxis to specific stimuli (*e.g. α*-methylasparate or nickel) and their combinations. The parameter values used in the model are consistent with those reported in Middlebrooks *et al*. (2021) where a different experimental system with spatially and temporally varying gradients was used to evaluate the multiscale model. Bacteria profiles were qualitatively similar to those published by Kalinin et al. (2010) for competing attractants; they set up two attractant gradients in opposite ends of a channel to compare the strength of responses to serine and *α*-methylasparate. From their work they concluded that the chemotactic response of *E. coli* depended on the number ratio of chemotaxis receptor proteins Tar/Tsr. In this work, we only considered cases where bacteria and chemoeffector sources were positioned on opposite sides of the channel, which means that two chemoeffectors were mixed first and then injected into one arm of the top layer of our device. One advantage to this approach is that the bacterial suspension was not exposed to one of the chemoeffectors in the channel prior to the experimental observation period. Upon exposure to high concentrations of chemoeffectors bacteria adapt to reset their receptor binding affinity, so they are able to continue to respond to chemical gradients. The multiscale model used in our analysis does not include adaptation of the chemotactic response that occurs at the molecular level through methylation of the transmembrane chemoreceptors. This methylation reaction is to ensure that the receptor is reset to appropriate baseline levels if local chemoeffector concentrations increase significantly. In our experimental system, bacteria swim up and down a relatively shallow gradient and experience continuous and gradual changes in chemoeffector concentrations. Thus, CheA activation by receptor binding and CheR/CheB activities are able to balance out, eliminating the need to specify methylation kinetics in the model (Lele et al., 2015). In conclusion, our study showed that a multiscale mathematical model can be used to predict bacterial responses to chemical mixtures. It provides insights to tune bacteria chemotactic response by modulating the attractant/repellent concentration for controlling *E. coli* biofilm formation in agricultural, medical and environmental practices.

## Acknowledgements

We acknowledge the Keck Center for Cellular imaging for their help to acquire the images. Dr. Xiaopu Wang for discussing about microfluidic device, and members in Dr. James P. Landers’ lab for helping with plasma treatment. This research was made possible in part by a grant from The Gulf of Mexico Research Initiative.

## Supporting Information

Additional supporting information may be found online in the Supporting Information section at the end of the article.

## Notes

### Competing Interest Statement

The authors have declared no competing interest.

## References

Aburto, A., & Peimbert, M. (2011). Degradation of a benzene–toluene mixture by hydrocarbon-adapted bacterial communities. Annals of microbiology, 61(3), 553–562.

Anderson, T. A., Guthrie, E. A., & Walton, B. T. (1993). Bioremediation in the rhizosphere. Environmental Science & Technology, 27(13), 2630–2636.

Babalola, O. O. (2010). Beneficial bacteria of agricultural importance. Biotechnology letters, 32(11), 1559–1570.

Berg, H. C., Brown, D. A. (1972). Chemotaxis in *Escherichia coli* analysed by three-dimensional tracking. Nature, 239(5374), 500–504.

Berg, H. C., Turner, L. (1990). Chemotaxis of bacteria in glass capillary arrays. *Escherichia coli*, motility, microchannel plate, and light scattering. Biophysical Journal, 58(4), 919–930.

Bergman K. (1990). The role of bacterial chemotaxis in initiation of rhizobium legume symbiosis for nitrogen-fixation. Journal of Chemical Ecology, 16(1), 114–115.

Clarke, S., & Koshland, D. E. (1979). Membrane receptors for aspartate and serine in bacterial chemotaxis. Journal of Biological Chemistry, 254(19), 9695–9702.

Feng, P., Ye, Z., Kakade, A., Virk, A. K., Li, X., & Liu, P. (2019). A review on gut remediation of selected environmental contaminants: possible roles of probiotics and gut microbiota. Nutrients, 11(1), 22.

Flint, H. J., Scott, K. P., Louis, P., & Duncan, S. H. (2012). The role of the gut microbiota in nutrition and health. Nature reviews Gastroenterology & hepatology, 9(10), 577–589.

Grebe, T. W., & Stock, J. (1998). Bacterial chemotaxis: the five sensors of a bacterium. Current Biology, 8(5), R154–R157.

Jasuja, R., Trentham, D. R., & Khan, S. (1999). Response tuning in bacterial chemotaxis. Proceedings of the National Academy of Sciences, 96(20), 11346–11351.

Kalinin, Y., Neumann, S., Sourjik, V., & Wu, M. (2010). Responses of *Escherichia coli* bacteria to two opposing chemoattractant gradients depend on the chemoreceptor ratio. Journal of Bacteriology, 192(7), 1796–1800.

Keilberg, D., & Ottemann, K. M. (2016). How *Helicobacter pylori* senses, targets and interacts with the gastric epithelium. Environmental microbiology, 18(3), 791–806.

Lele, P. P., Shrivastava, A., Roland, T., & Berg, H. C. (2015). Response thresholds in bacterial chemotaxis. Science advances, 1(9), e1500299.

Meckenstock, R. U., Elsner, M., Griebler, C., Lueders, T., Stumpp, C., Aamand, J., … & van Breukelen, B. M. (2015). Biodegradation: updating the concepts of control for microbial cleanup in contaminated aquifers. Environmental science & technology, 49(12), 7073–7081.

Middlebrooks, S. A. (1993). The chemotactic response of Escherichia coli to combined repellent and attractant stimuli. (M.S. Thesis) University of Virginia.

Middlebrooks, S. A., Zhao, X., Ford, R. M., & Cummings, P. T. (2021). A mathematical model for Escherichia coli chemotaxis to competing stimuli. Biotechnology and Bioengineering, 118(12), 4678–4686.

Mowbray, S. L., & Koshland, D. E. (1987). Additive and independent responses in a single receptor: Aspartate and maltose stimuli on the tar protein. Cell, 50(2), 171–180.

Pandey, G., & Jain, R. K. (2002). Bacterial chemotaxis toward environmental pollutants: role in bioremediation. Appl. Environ. Microbiol., 68(12), 5789–5795.

Park, J., Aminzare, Z. A. (2022). Mathematical Description of Bacterial Chemotaxis in Response to Two Stimuli. Bull. Math. Biol., 84(9).

Rivero, M. A., Tranquillo, R. T., Buettner, H. M., & Lauffenburger, D. A. (1989). Transport models for chemotactic cell populations based on individual cell behavior. Chemical Engineering Science, 44(12), 2881–2897.

Sourjik, V. (2004). Receptor clustering and signal processing in E. coli chemotaxis. Trends in microbiology, 12(12), 569–576.

Sourjik, V., Armitage, J. P. (2010). Spatial organization in bacterial chemotaxis. The EMBO journal, 29(16), 2724–2733.

Strauss, I., Frymier, P. D., Hahn, C. M., & Ford, R. M. (1995). Analysis of Bacterial Migration. II. Studies with Multiple Attractant Gradients. AICHE Journal, 41(2), 402–414.

Tsiaoussis, J., Antoniou, M. N., Koliarakis, I., Mesnage, R., Vardavas, C. I., Izotov, B. N., … & Tsatsakis, A. (2019). Effects of single and combined toxic exposures on the gut microbiome: current knowledge and future directions. Toxicology letters, 312, 72–97.

Tso, W. W., Adler, J. (1974). Negative chemotaxis in *Escherichia coli*. Journal of bacteriology, 118(2), 560–576.

Wang, X. (2013). Application of Microfluidic Devices for the Study of Bacterial Chemotaxis to NAPL Components in Porous Media. (Ph.D. Dissertation) University of Virginia.

Wang, X., Atencia, J., Ford, R. M. (2015). Quantitative Analysis of Chemotaxis Towards Toluene by *Pseudomonas putida* in a convection-free device. Biotechnology and Bioengineering, 112(5), 896–904.

Wolfe AJ, Conley MP, Kramer TJ, Berg HC. (1987). Reconstitution of signaling in bacterial chemotaxis. Journal of Bacteriology, 169, 1878–1885.

Worku, M. L., Sidebotham, R. L., Walker, M. M., Keshavarz, T., Karim, Q. N. (1999). The relationship between *Helicobacter pylori* motility, morphology and phase of growth: implications for gastric colonization and pathology. Microbiology, 145(10), 2803–2811.

Yawata, Y., Cordero, O. X., Menolascina, F., Hehemann, J. H., Polz, M. F., & Stocker, R. (2014). Competition-dispersal tradeoff ecologically differentiates recently speciated marine bacterioplankton populations. Proceedings of the National Academy of Sciences, 111(15), 5622–5627.

Zhang, X., Si, G., Dong, Y., Chen, K., Ouyang, Q., Luo, C., & Tu, Y. (2019). Escape band in Escherichia coli chemotaxis in opposing attractant and nutrient gradients. Proceedings of the National Academy of Sciences, 116(6), 2253–2258.

