## Supplementary Information for "*Escherichia coli* chemotaxis to competing stimuli in a microfluidic device with a constant gradient"

**Correspondence**

Supplementary Information

### 1. SI Mathematical Model

#### 1.1 Signal transduction kinetics

The autophosphorylation of histidine kinase when the ternary receptor complex (Tar-CheW-CheA, which we refer to as CheA, hereafter) is unbound is represented as

$$\text{CheA+ P} \overset{K_1}{\Leftrightarrow} \text{CheA-P} \tag{S1}$$

$$\text{CheA}_T = \text{CheA+ CheA-P} \tag{S2}$$

where $\text{CheA}_T$, a fixed value, is the summation of CheA and CheA-P, CheA is the unbound receptor complex, CheA-P is the phosphorylated receptor complex, and $K_1 = 1.8$ mM (Stewart, 2005) is the dissociation constant of phosphorylation of the unbound receptor complex.

We assume that the binding of α-methylaspartate (Asp) and nickel ion (Ni) to the receptor complex CheA can be regarded as a competitive adsorption equilibrium, represented as

$$\text{CheA + Asp} \overset{K_{dA}}{\Longleftrightarrow} \text{Asp-CheA} \tag{S3}$$

$$\text{CheA + Ni} \overset{K_{dN}}{\Longleftrightarrow} \text{Ni-CheA} \tag{S4}$$

$$\text{Ni-CheA + P} \overset{K_2}{\Leftrightarrow} \text{CheA- P} \tag{S5}$$

$$[\text{CheA}_T] = [\text{CheA}] + [\text{Asp-CheA}] + [\text{Ni-CheA}] + [\text{CheA- P}] + [\text{Ni-CheA- P}] \tag{S6}$$

In the above Equations (S3-S6), Asp-CheA and Ni-CheA represent bound receptor complexes of α-methylaspartate and nickel ions, $\text{CheA}_T$ is the total amount of receptor and is a fixed number, $K_{dA} = 0.64$ mM (Clarke and Koshland, 1979) is the dissociation constant of α-methylaspartate and $K_{dN} = 0.54$ mM (Middlebrooks, 1993) is the dissociation constant of nickel ions.

Two parameters were defined to represent the signaling transduction mechanism at the molecular scale. $\gamma = \frac{K_1}{[P]}$ is the signaling efficiency, and $\kappa = \frac{K_1}{K_2}$ is the repellent sensitivity coefficient.

We assume that the binding of aspartate (and its derivative α-methylaspartate) and nickel to the Tar receptor is competitive. We know that aspartate binds to the periplasmic domain of the Tar receptor (Park et al., 2011) and nickel does not require the periplasmic NikA binding protein for the chemotaxis reaction (Englert et al., 2010). However, the specific site on the Tar receptor where nickel binds is still debated in the literature. Englert *et al.* (2010) found that Ni binds specifically to the periplasmic domain of the Tar receptor, and that Ni uptake into the cell is not required for repellent taxis in response to Ni. However, Bi *et al.* (2018) reported that Ni is detected not by the periplasmic domain, but most likely by the HAMP domain (in the cytoplasm region) and its junction with the transmembrane domain of the Tar receptor. The reason for their conclusion is that they observed that a Tar mutant that completely lacks the periplasmic domain still showed a repellent response to Ni.

After binding with aspartate, the Tar receptor undergoes conformational changes, including a piston motion in the periplasmic domain and a bending motion in the transmembrane domain. These conformational changes will partially hinder the binding of nickel. However, the mechanism of nickel binding is not fully understood. There is also the possibility that aspartate and nickel can bind to the Tar receptor at the same time. Gardina *et al.* (1998) suggested that the binding of aspartate and maltose can be non-competitive if the ligands are restricted to a particular orientation.

#### 1.2 Tumbling probability

Berg and Brown (1972) observed that the logarithm of mean run time increased with respect to the time rate of change in the number of bound receptors according to

$$\ln\,\tau = \ln\,\tau_0 + \alpha\,\frac{dN_b}{dt} \tag{S7}$$

where $\tau$ is the mean run time, $\tau_0$ is the mean run time in absence of the chemoattractant gradient, $\alpha$ is the proportionality constant, and $N_b$ is the number of bound receptors in the presence of chemoattractant.

In our model, we replaced the number of bound receptors with the amount of signaling complex CheA-P, which is the step where information from nickel and α-methylaspartate is integrated. The mean run time is the reciprocal of the tumbling probability $\tau = 1/p_t$, and after substitution into Equation (S7), we get

$$\frac{p_t^{+/-}}{p_0} = \exp\left(-\varepsilon \frac{D}{Dt}\frac{[CheA-P]}{[CheA-P]_0}\right) \quad \text{(S8)}$$

where $p_0$ is the tumbling probability in the absence of a chemoattractant gradient, $\varepsilon$ is a proportionality constant, $[CheA - P]$ is the amount of signaling complex in the presence of chemoeffectors, and $[CheA - P]_0$ is the amount of signaling complex in the absence of chemoeffectors.

Furthermore, the chemotactic velocity of a bacteria population is defined in terms of individual cell properties as (Chen et al., 1998)

$$V_c = \frac{2}{3} v \frac{p^- - p^+}{p^- + p^+} \quad \text{(S9)}$$

where $p^+$ is the probability per unit time that a cell moving up the gradient will tumble and change direction to join the down-gradient population of bacteria, $p^-$ $is$ the probability per unit time that a cell moving down the gradient will tumble and change direction to join the up-gradient population of bacteria and *v* is the bacterium swimming speed; however *v* is reduced to $\frac{1}{2}v$ if we project the velocity from three-dimensional to one-dimensional, therefore, $V_c = \frac{1}{3} v \frac{p^- - p^+}{p^- + p^+}$.

### 1.3. Derivation of the chemotactic velocity under various conditions

1.3.1 Phosphorylation of histidine kinase in the absence of chemoeffectors

From Equation (S1), we have

$$K_1 = \frac{[CheA][P]}{[CheA-P]} \quad \text{(S10)}$$

and we know that $[CheA] = [CheA_T] - [CheA - P]$ from Equation (S2). Therefore, we can get

$$[CheA - P] = \frac{[CheA_T][P]}{K_1 + [P]} = \frac{[CheA_T]}{\frac{K_1}{[P]} + 1} = \frac{[CheA_T]}{\gamma + 1} \quad \text{(S11)}$$

where $\gamma = \frac{K_1}{P}$, is the signaling efficiency.

1.3.2 Chemotactic velocity in presence of attractant

From Equation (S3), we know that $K_{dA} = \frac{[CheA][Asp]}{[CheA-Asp]}$ (S12) and we can rearrange that expression to get $[CheA - Asp] = \frac{[CheA][Asp]}{K_{dA}}$ (S13)

The total amount of $[CheA]$ is now

$$[CheA_T] = [CheA] + [CheA - P] + [CheA - Asp] \quad \text{(S14)}$$

Substitution of Equations (S12-S13) into (S14) gives

$$[CheA] = \frac{[CheA_T]}{1+\frac{[P]}{K_1}+\frac{[Asp]}{K_{dA}}} \quad \text{(S15)}$$

Then substitution of Equation (S15) into a rearrangement of Equation (S10) gives

$$[CheA - P] = \frac{[CheA_T][P]}{K_1+[P]+K_1\frac{[Asp]}{K_{dA}}} = \frac{[CheA_T]}{1+\frac{K_1}{[P]}+\frac{K_1[Asp]}{[P]K_{dA}}} = \frac{[CheA_T]}{1+\gamma+\gamma\frac{[Asp]}{K_{dA}}} \quad \text{(S16)}$$

where $\gamma = \frac{K_1}{P_i}$ is the signaling efficiency and $K_{dA}$ is the dissociation constant for aspartate binding.

From Equation (S7), $\frac{DN_b^{+/-}}{Dt} = \frac{\partial N_b}{\partial t} \pm v\frac{\partial N_b}{\partial x}$, and we use $[CheA - P]$ to replace the number of bound receptors $N_b$, which gives

$$p_t^{+/-} = p_0 \exp\left[-\alpha\left(\frac{\partial[CheA-P]}{\partial t} \pm v\frac{\partial[CheA-P]}{\partial x}\right)\right] \quad \text{(S17)}$$

We can also write Equation (S17) as $\frac{p_t^{+/-}}{p_0} = \exp\left(-\sigma\left(\frac{\partial}{\partial t}(\frac{[CheA-P]}{[CheA-P]_0}) \pm v\frac{\partial}{\partial x}(\frac{[CheA-P]}{[CheA-P]_0})\right)\right)$, where $\sigma$ is the stimuli sensitivity coefficient. The expression for $[CheA - P]$ is a function of $[Asp]$, and $[Asp]$ is independent of time under steady state conditions, so Equation (S17) can be simplified to $p_t^{+/-} = p_0 \exp\left[\mp\sigma\left(v\frac{\partial}{\partial x}(\frac{[CheA-P]}{[CheA-P]_0})\right)\right]$ (S18)

Substituting the above equation into Equation (S9), yields the chemotactic velocity of bacteria in the presence of attractant

$$V_c = \frac{1}{3}v\frac{e^{\sigma v\frac{\partial}{\partial x}\frac{[CheA-P]}{[CheA-P]_0}} - e^{+\sigma v\frac{\partial}{\partial x}\frac{[CheA-P]}{[CheA-P]_0}}}{e^{\sigma v\frac{\partial}{\partial x}\frac{[CheA-P]}{[CheA-P]_0}} - e^{+\sigma v\frac{\partial}{\partial x}\frac{[CheA-P]}{[CheA-P]_0}}} = v\tanh(\sigma v\frac{\partial}{\partial x}\frac{[CheA-P]}{[CheA-P]_0}) \sim v\sigma v\frac{\partial}{\partial x}\frac{[CheA-P]}{[CheA-P]_0}$$

when $\sigma v\frac{\partial}{\partial x}(\frac{[CheA-P]}{[CheA-P]_0})$ is small. Therefore,

$$V_c = \frac{1}{3}\sigma v^2\frac{\partial}{\partial x}\left(\frac{1+\gamma}{1+\gamma+\gamma\frac{[Asp]}{K_{dA}}}\right) = \frac{1}{3}\sigma v^2\frac{(\frac{\gamma+1}{\gamma})K_{dA}}{((\frac{\gamma+1}{\gamma})K_{dA}+[Asp])^2}\frac{\partial([Asp])}{\partial x} \quad \text{(S19)}$$

1.3.3 Chemotactic velocity in the presence of repellent

Receptors bound to nickel are able to be phosphorylated as shown in Equation (S5), therefore,

$$[CheA_T] = [CheA] + [CheA-P] + [Ni-CheA] + [Ni-CheA-P] \quad \text{(S20)}$$

From Equation (S1), we know that $[CheA-P] = \frac{[CheA][P]}{K_1}$ (S21)

From Equation (S4), we can get

$K_{dN} = \frac{[Ni][CheA]}{[Ni-CheA]}$, so $[Ni-CheA] = \frac{[Ni][CheA]}{K_{dN}}$ (S22)

From Equation (S5), we know that

$K_2 = \frac{[Ni-CheA][P]}{[Ni-CheA-P]}$, so $[Ni-CheA-P] = \frac{[Ni-CheA][P]}{K_2}$ (S23)

Substitution of Equations (S21-S23) into Equation (S20) gives $[CheA] = \frac{[CheA_T]}{1+\frac{[P]}{K_1}+(1+\frac{[P]}{K_2})\frac{[Ni]}{K_{dN}}}$

Therefore $[CheA-P] = \frac{[CheA_T]}{1+\frac{K_1}{[P]}+\frac{[Ni]\,K_1}{K_{dN}[P]}+\frac{K_1\,[Ni]}{K_2K_{dN}}}$ and

$$[Ni-CheA-P] = \frac{[CheA_T]}{1+\frac{[P]}{K_1}+(1+\frac{[P]}{K_2})\frac{[Ni]}{K_{dN}}}\frac{[Ni][P]}{K_{dN}K_2} = \kappa[CheA-P]\frac{[Ni]}{K_{dN}}$$

Thus, $[CheA-P]_{total} = (1+\kappa\frac{[Ni]}{K_{dN}})[CheA-P] = (1+\kappa\frac{[Ni]}{K_{dN}})\frac{[CheA_T]}{1+\frac{K_1}{[P]}+\frac{[Ni]\,K_1}{K_{dN}[P]}+\frac{K_1\,[Ni]}{K_2K_{dN}}} = (1+$
$\kappa\frac{[Ni]}{K_{dN}})\frac{[CheA_T]}{1+\gamma+(k+\gamma)\frac{[Ni]}{K_{dN}}}$

where $\gamma = \frac{K_1}{[P]}$, is the signaling efficiency, $\kappa = \frac{K_1}{K_2}$ is the repellent sensitivity coefficient, $K_{dN}$ is the dissociation constant for nickel ion binding.

Substitution of $[CheA-P]_{total}$ into Equation (S18), and then into Equation (S9) gives the chemotactic velocity in the presence of repellent,

$$V_c = \frac{1}{3}\sigma v^2\frac{\partial}{\partial x}(\frac{[CheA-P]}{[CheA-P]_0}) = \frac{1}{3}\sigma v^2\frac{\partial}{\partial x}(\frac{(1+\gamma)\left(1+\kappa\frac{[Ni]}{K_{dN}}\right)}{1+\gamma+(k+\gamma)\frac{[Ni]}{K_{dN}}}) = -\frac{1}{3}\sigma(\kappa-1)v^2\frac{(\frac{\gamma+1}{\gamma})K_{dN}}{[\left(\frac{\gamma+1}{\gamma}\right)K_{dN}+(\frac{\kappa+\gamma}{\gamma})[Ni]]^2}\frac{\partial([Ni])}{\partial x}$$

(S24)

1.3.4 Chemotactic velocity in the presence of combined chemoeffectors

When both stimuli are present, the total amount of receptor complexes can be represented as

$$[CheA_T] = [CheA] + [CheA-P] + [Asp-CheA] + [Ni-CheA] + [Ni-CheA-P] \quad \text{(S25)}$$

Substitution of Equations (S13) and (S21-S23) into Equation (S25) yields

$$[CheA] = \frac{[CheA_T]}{1+\frac{[P]}{K_1}+\frac{[Asp]}{K_{dA}}(1+\frac{[P]}{K_2})\frac{[Ni]}{K_{dN}}}$$

Therefore $[CheA-P]=\frac{[CheA_T]}{1+\frac{K_1}{[P]}+\frac{[Asp]K_1}{K_{dA}[P]}+(\frac{K_1}{[P]}+\frac{K_1}{K_2})\frac{[Ni]}{K_{dN}}}$ and

$$[Ni-CheA-P]=\frac{[CheA_T]}{1+\frac{K_1}{[P]}+\frac{[Asp]}{K_{dA}}\frac{K_1}{[P]}+(\frac{K_1}{[P]}+\frac{K_1}{K_2})\frac{[Ni]}{K_{dN}}}\frac{[Ni][P]}{K_{dN}K_2}=\kappa[CheA-P]\frac{[Ni]}{K_{dN}}$$

Thus $[CheA-P]_{total}=\left(1+\kappa\frac{[Ni]}{K_{dN}}\right)[CheA-P]=(1+\kappa\frac{[Ni]}{K_{dN}})\frac{[CheA_T]}{1+\frac{K_1}{[P]}+\frac{[Asp]K_1}{K_{dA}[P]}+(\frac{K_1}{[P]}+\frac{K_1}{K_2})\frac{[Ni]}{K_{dN}}}=(1+\kappa\frac{[Ni]}{K_{dN}})\frac{[CheA_T]}{1+\gamma+\gamma\frac{[Asp]}{K_{dA}}+(k+\gamma)\frac{[Ni]}{K_{dN}}}$

where $\gamma=\frac{K_1}{[P]}$, is the signaling efficiency, $\kappa=\frac{K_1}{K_2}$ is the repellent sensitivity coefficient, $K_{dA}$ is the dissociation constant for aspartate ion binding, and $K_{dN}$ is the dissociation constant for nickel ion binding.

Substitution of $[CheA-P]_{total}$ into Equation (S18), and then Equation (S9) gives the chemotactic velocity in the presence of combined chemoeffectors,

$$V_c=\frac{1}{3}\sigma v^2\frac{\partial}{\partial x}(\frac{[CheA-P]}{[CheA-P]_0})=\frac{1}{3}\frac{\partial}{\partial x}(\frac{(1+\gamma)\left(1+\kappa\frac{[Ni]}{K_{dN}}\right)}{1+\gamma+\gamma\frac{[Asp]}{K_{dA}}+(k+\gamma)\frac{[Ni]}{K_{dN}}})=\frac{1}{3}\frac{\sigma v^2(1+\gamma)\gamma}{\left[1+\gamma+\gamma\frac{[Asp]}{K_{dA}}+(k+\gamma)\frac{[Ni]}{K_{dN}}\right]^2}\{(1+\kappa\frac{[Ni]}{K_{dN}})\frac{\partial}{\partial y}(\frac{[Asp]}{K_{dA}})-(\kappa-1+\kappa\frac{[Asp]}{K_{dA}})\frac{\partial}{\partial y}(\frac{[Ni]}{K_{dN}})\} \quad \text{(S26)}$$

### 2. SI Experimental Methods

#### 2.1 Microfluidic device fabrication – step to create PDMS components

A PDMS solution was made using a Sylgard 184 silicone elastomer kit (Dow Corning Corporation, MI). The base and curing agent were mixed and the resulting solution was cured on top of the silicon mold with the channel designs. Then, the Y-shaped channel that makes up the top layer and the four cross channels that make up the bottom layer were cut in rectangular blocks from the silicon mold and peeled off. Both of the layers were subsequently treated with a plasma cleaner (PlasmaEtch, NV) to increase the hydrophilicity of PDMS.

### 3. SI Results and Discussion

#### 3.1 Validation of diffusion in the microfluidic device

An experiment was conducted to confirm convection-free cross channels and test the accuracy of the microfluidic device. A source of fluorescent uranine was provided via one arm of the upper Y-shaped channel and buffer via the other arm, which then served as a sink for uranine in the lower cross channels. Microscopic images of an underlying cross channel taken at several times are shown in Figure S1. The gradually increasing penetration of fluorescence intensity into the cross channel was qualitatively consistent with a diffusive process. Furthermore, the experimental observations and theoretical predictions with $D = 0.49 \times 10^{-9}$ m$^2$/s (cite reference) match well. This result confirms that diffusion was the governing process in the cross channel of this microfluidic device. Note the experimental data in Figure S1 show fluorescence intensity values only in the range from 0.45 to 1.5 mm because the intensity close to the vias was too high and saturated the camera detection limits.

The Einstein relation was used to estimate the time required to reach steady state in the cross channels:

$$L^2 = 2Dt \qquad \text{(S27)}$$

where $L$ is the channel length of 1.5 mm and $D = 0.49 \times 10^{-9}$ m$^2$/s is the diffusion coefficient for uranine. Solving Equation (S27) yielded t = 38 min as the time needed to reach the steady state in the channel.

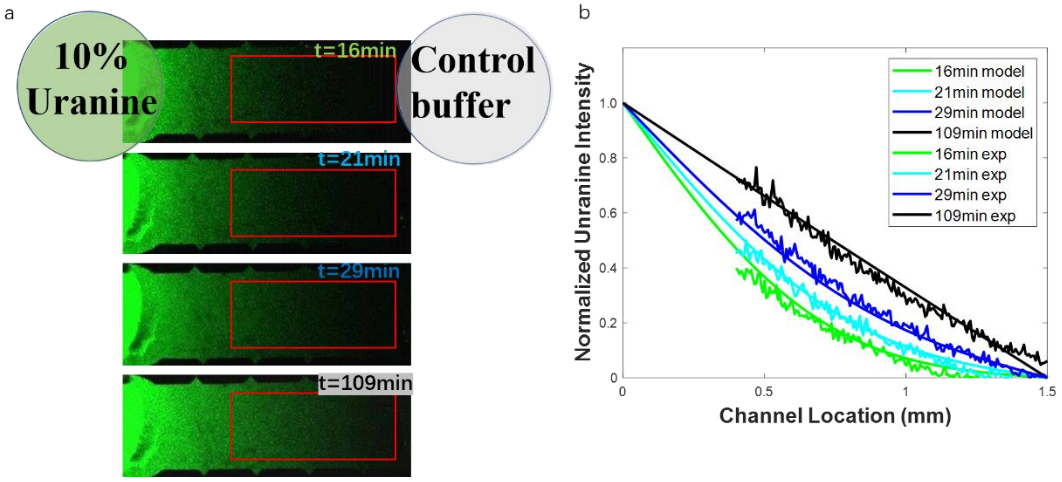


Figure S1. (a) 10% uranine distribution in a cross channel at several time points. There is a constant source of 10% uranine on the left-hand side of the channel and a constant sink on the right-hand side; the fluorescence intensity is proportional to uranine concentration. Red rectangular box indicates the region over which data was analyzed. (b) Normalized uranine intensity along the channel (scattered data) and the model equation results (solid lines) are compared at several times.

#### 3.2 *E. coli* random motility coefficient

For diffusive processes the Boltzmann transformation variable can be used to collapse data collected at different times onto a single spatial profile. The random motility coefficient can then be estimated from the following arrangement of the transformed bacterial diffusion equation:

$$\mu_0(\varepsilon) = \frac{-\int_{y(\infty)}^{y(\varepsilon)} \varepsilon\, dy}{2\,(\frac{dy}{d\varepsilon})_\varepsilon}$$

where y is the normalized bacteria intensity and $\varepsilon$ is the Boltzmann transformation variable. The numerator $\int_{y(\infty)}^{y(\varepsilon)} \varepsilon\, dy$ is the area under the curve, while the denominator contains the slope of the curve. Applying this analysis to the curve fit for Model 1 in the inset of Figure 3 yields

$$\mu_0(0) = \frac{-\int_{y(\infty)}^{y(0)} \varepsilon\, dy}{2\,(\frac{dy}{d\varepsilon})_0} = \frac{-0.0129}{2\times(-49.5)} = 0.00013\, mm^2/s = 1.3\times 10^{-10}\, m^2/s$$

#### 3.3 Derivation of signaling efficiency $\gamma$ from random motility coefficient $\mu_0$

The averaged random motility coefficient determined from this study was $1.3 \pm 0.21 \times 10^{-10}\, m^2/s$. This population-scale transport property can be related to the tumble probability, a cellular-scale parameter, according to Equation (4.4) from Chen et al. (1998),

$$\mu_0 = \frac{v^2}{3p_t} \qquad \text{(S28)}$$

where $v$ is bacteria swimming speed and $p_t$ is the tumble probability. For simplicity, the directional persistence term was not included in this analysis.

To calculate $p_t$, the value for *E. coli* swimming speed $v$ was substituted into Equation (S28). Reported values for swimming speed range from 10-30 $\mu m/s$ (Rivero et al., 1989); we chose $v = 15\ \mu m/s$. Then $p_t = 0.58\ s^{-1}$. Tumble probability $p_t$ can be further related to the switching frequency, which is the number of times that a flagella motor switched its direction of rotation divided by the duration of the recording. Since it takes two switches from CCW to CW and CW to CCW in order to tumble once, the switching frequency in Cluzel et al. (2000) is twice the tumbling probability we used in our model. Therefore, the switching frequency is $2p_t = 1.16\ s^{-1}$, which corresponds to a CheY-P concentration of 2.7 $\mu$M in Figure S2.

Recall that $[CheA - P] = \frac{[CheA_T]}{\gamma+1}$ in Equation (S11). And the transfer of phosphate to the response regulator CheY is represented as

$$\text{CheA-P} \xrightarrow{k} \text{CheY-P} \qquad \text{(S29)}$$

where CheY-P is the phosphorylated CheY and $k$ is the reaction rate constant of the phosphate transfer reaction.

We then define the concentration of phosphorylated CheY in the absence of chemoattractant as

$$[\mathrm{CheY-P}]_0 = \frac{\alpha CheA_T}{\gamma+1} \quad \text{(S30)}$$

where $\alpha = 1$ is a stoichiometric coefficient, $\mathrm{CheA_T}$ is the total amount of receptor, a fixed number $CheA_T = 7.9\ \mu\mathrm{M}$ (Edgington and Tindall, 2015), and $\gamma = \frac{K_1}{[P]}$ is the signaling efficiency.

Rearranging Equation (S30) to solve for the signaling efficiency

$$\gamma = \frac{\alpha CheA_T}{[\mathrm{CheY-P}]_0} - 1$$

and assuming a stoichiometric coefficient of one yields a value of $\gamma = 2$. A greater value for total amount of receptor $\mathrm{CheA_T}$ =13 μM is needed to match the experimentally-fitted value of $\gamma = 4$.

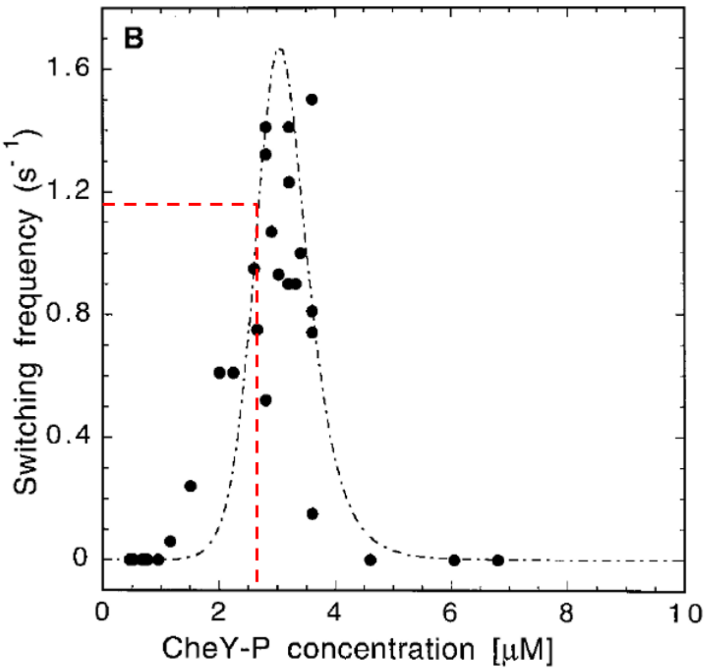


Figure S2. Switching frequency as a function of CheY-P concentration. Adapted from Cluzel et al. (2000).

#### 3.4 Modulating *E. coli* response to a chemoeffector with increasing concentrations of an opposing stimulus

In this section microscopic images, bacterial intensity profiles and model predictions at different concentration combinations are presented. Figure S3 shows that increasing Ni concentrations counter the attraction of α-methylaspartate and change the bacterial response from overall "attraction" to "repulsion". Figure S4 shows that increasing concentrations of an attractant counter the repellent effect of nickel; the combined chemotactic response to attractant and repellent almost cancels out each other at higher concentrations of the opposing stimulus.

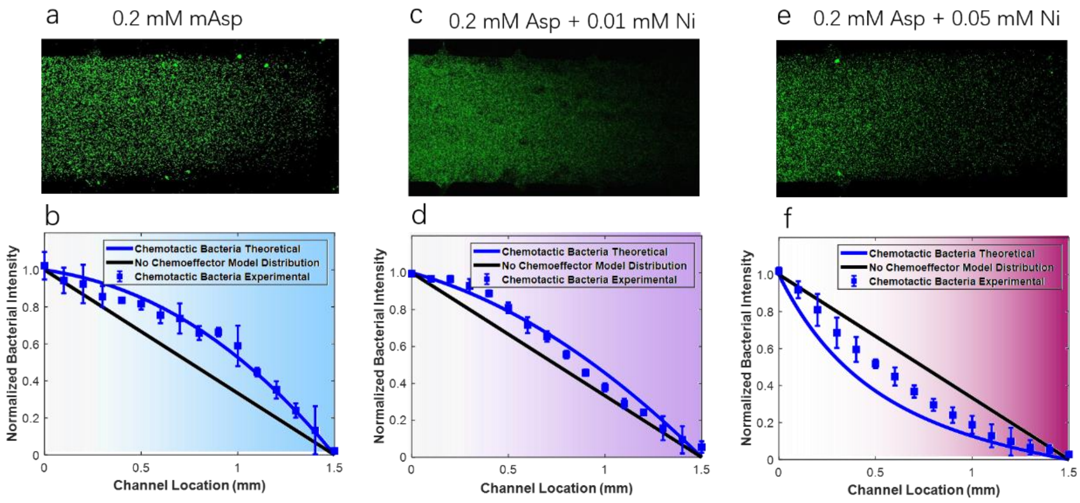


Figure S3. GFP-labeled *E. coli* distribution in a cross channel at steady state. There is a constant source of bacteria on the left-hand side of the channel, and a constant source of chemoeffector on the right-hand side: (a) 0.2 mM $\alpha$-methyl aspartate, (c) 0.2 mM $\alpha$-methyl aspartate and 0.01 mM nickel ions, and (e) 0.2 mM $\alpha$-methyl aspartate and 0.05 mM nickel ions. The fluorescence intensity is proportional to bacterial concentration and is plotted as normalized intensity as a function of location for bacteria response to (b) 0.2 mM $\alpha$-methyl aspartate, (d) 0.2 mM $\alpha$-methyl aspartate and 0.01 mM nickel ions, and (f) 0.2 mM $\alpha$-methyl aspartate and 0.05 mM nickel ions. Note that although the bacteria concentrations in the images are not the same, the gray level value in each image is calibrated itself using the approach in Methods section.

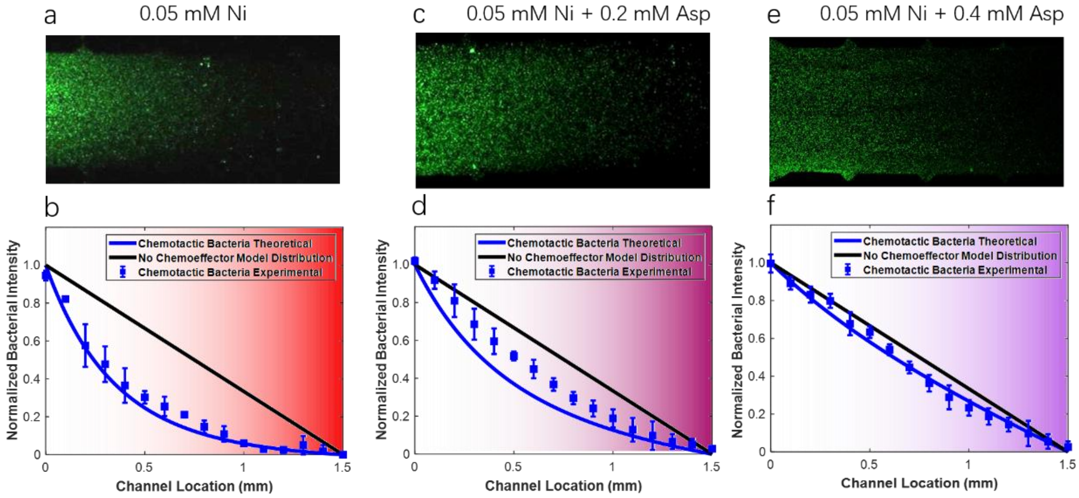


Figure S4. GFP-labeled *E. coli* distribution in a cross channel at steady state for a constant source of bacteria on the left-hand side of the channel and a constant source of chemoeffector on the right-hand side with (a) 0.05 mM nickel ions (c) 0.05 mM nickel ions and 0.2 mM $\alpha$-methyl aspartate and (e) 0.05 mM nickel ions and 0.4 mM $\alpha$-methylaspartate. The fluorescence intensity is proportional to bacterial concentration and is plotted as normalized intensity as a function of location for bacteria response to (b) 0.05 mM nickel ions, (d) 0.05 mM nickel ions and 0.2 mM $\alpha$-methyl aspartate, and (f) 0.05 mM nickel ions and 0.4 mM $\alpha$-methyl aspartate. Note that although the bacteria concentrations in the images are not the same, the gray level value in each image is calibrated to itself using the approach documented in the Methods section.

**References**

Berg, H. C., Brown, D. A. (1972). Chemotaxis in *Escherichia coli* analysed by three-dimensional tracking. *Nature*, *239*(5374), 500-504.

Bi, S., Jin, F., & Sourjik, V. (2018). Inverted signaling by bacterial chemotaxis receptors. *Nature communications*, *9*(1), 1-13.

Borkovich, K. A., & Simon, M. I. (1990). The dynamics of protein phosphorylation in bacterial chemotaxis. *Cell*, *63*(6), 1339-1348.

Chen, K. C., Cummings, P. T., & Ford, R. M. (1998). Perturbation expansion of Alt's cell balance equations reduces to Segel's one-dimensional equations for shallow chemoattractant gradients. SIAM Journal on Applied Mathematics, *59*(1), 35-57.

Clarke, S., & Koshland, D. E. (1979). Membrane receptors for aspartate and serine in bacterial chemotaxis. *Journal of Biological Chemistry*, *254*(19), 9695-9702.

Cluzel, P., Surette, M., & Leibler, S. (2000). An ultrasensitive bacterial motor revealed by monitoring signaling proteins in single cells. *Science*, *287*(5458), 1652-1655.

Edgington, M. P., & Tindall, M. J. (2015). Understanding the link between single cell and population scale responses of *Escherichia coli* in differing ligand gradients. *Computational and structural biotechnology journal*, *13*, 528-538.

Englert, D. L., Adase, C. A., Jayaraman, A., & Manson, M. D. (2010). Repellent taxis in response to nickel ion requires neither $Ni^{2+}$ transport nor the periplasmic NikA binding protein. *Journal of bacteriology*, *192*(10), 2633-2637.

Gardina, P. J., Bormans, A. F., & Manson, M. D. (1998). A mechanism for simultaneous sensing of aspartate and maltose by the Tar chemoreceptor of *Escherichia coli*. *Molecular microbiology*, *29*(5), 1147-1154.

Middlebrooks, S. A., Zhao, X., Ford, R. M., & Cummings, P. T. (2021). A mathematical model for Escherichia coli chemotaxis to competing stimuli. *Biotechnology and Bioengineering, 118*(12), 4678-4686.

Park, H., Im, W., & Seok, C. (2011). Transmembrane signaling of chemotaxis receptor tar: insights from molecular dynamics simulation studies. *Biophysical journal, 100*(12), 2955-2963.

Rivero, M. A., Tranquillo, R. T., Buettner, H. M., & Lauffenburger, D. A. (1989). Transport models for chemotactic cell populations based on individual cell behavior. *Chemical Engineering Science*, *44*(12), 2881-2897.

Stewart, R. C. (2005). Analysis of ATP binding to CheA containing tryptophan substitutions near the active site. *Biochemistry*, *44*(11), 4375-4385.
